# Distribution of antibiotic resistance genes across contrasted tropical agroecosystems in Réunion Island

**DOI:** 10.64898/2026.01.27.702181

**Authors:** Rieux Adrien, Dolivet-Maréchal Marion, Doizy Anna, Chiroleu Frederic, Mottes Charles, Soti Valérie, Darnaudery Marie, Matthieu N. Bravin, Cardinale Eric, Baldet Thierry, Doelsch Emmanuel

## Abstract

This study presents the first exploratory assessment of antibiotic resistance genes (ARGs) and antibiotic residues in agricultural environments on Réunion Island, a French tropical territory in the south-western Indian Ocean. Sixteen samples from diverse matrices (manure, soil, water, and vegetables) across different agroecosystems were analyzed using high-throughput qPCR targeting 332 ARGs and chemical methods targeting 58 antibiotic compounds and trace elements. ARGs were widely detected across all matrices, with highest abundance observed in amended soils and manure. Surprisingly, ARG profiles, in terms of both abundance and average number, were comparable between unamended soils and natural soils. Antibiotic residues were found in only five manure and soil samples, with no clear correlation between the presence of these residues, trace elements and ARG abundance. Organic amendments significantly increased ARG levels in soils and non-metric multidimensional scaling revealed that ARG profiles clustered primarily by matrix type rather than by location. High-risk ARGs were widely prevalent, with 86% detected and 23% ubiquitous across all samples, and their occurrence in water and raw vegetables suggests potential human exposure through the food chain. This study highlights the influence of agricultural practices on environmental antimicrobial resistance in tropical island contexts and supports the need for expanded One Health surveillance integrating the environmental, animal and human compartments.

**Synopsis:** This study shows that agricultural practices can shape the environmental spread of antibiotic resistance in tropical island ecosystems, highlighting potential risks for ecosystems, food safety, and human health.

**Graphical abstract:** 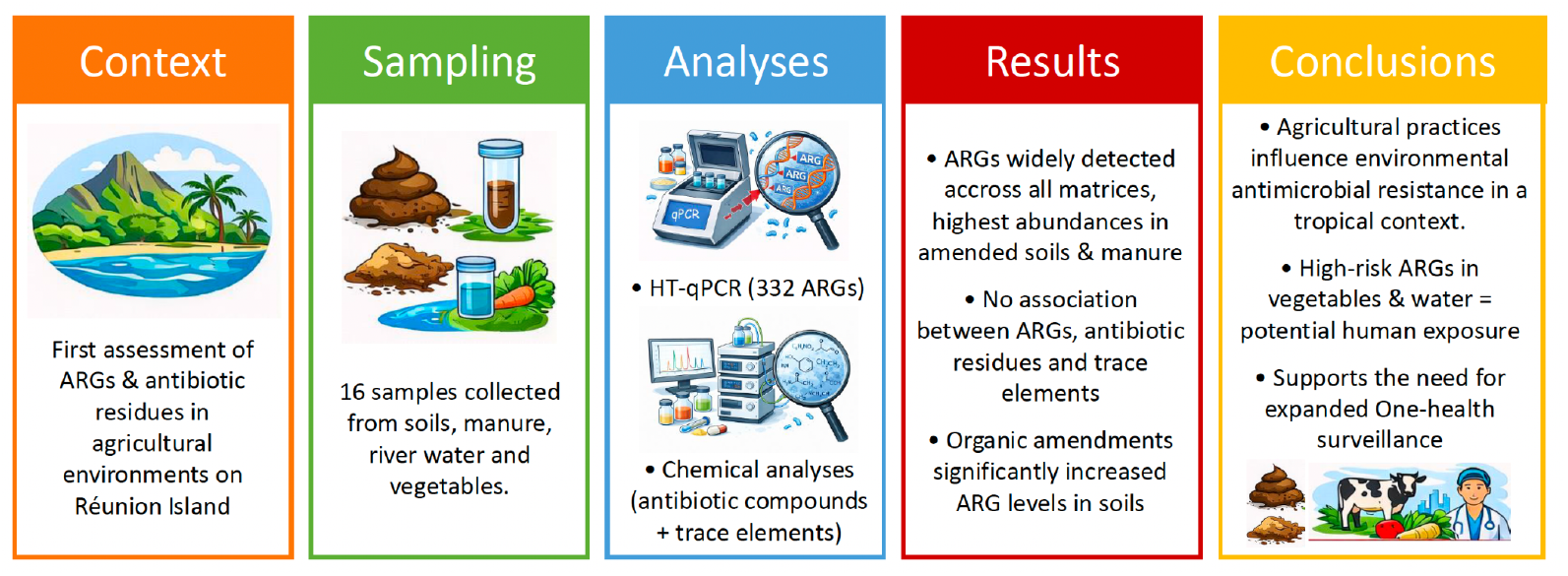

## Introduction

Antimicrobial resistance (AMR) is one of the biggest threats to global health today^1^. Although AMR is initially a natural phenomenon shaped by the long evolutionary arms race between microorganisms over millennia^2^, its importance has increased significantly in recent decades due to human practices such as the intensive and often inappropriate use of antibiotics in both human and animal health^3,4^. This has led to the emergence of multidrug-resistant “superbugs” capable of evading almost all standard antibiotic treatments and jeopardizing the effectiveness of even last-resort therapies^3^. Beyond the clinical and therapeutic challenges in public and veterinary health, the environmental dimension of AMR is attracting increasing attention^5,6^. Antibiotics used in medicine and animal husbandry frequently enter agroecosystems through poor wastewater and manure management and consequently enter natural ecosystems through agricultural runoff and leaching, along with antibiotic resistant bacteria (ARB) carrying antibiotic resistance genes (ARGs)^5^. Consequently, ARB and ARGs have now been identified across a wide range of natural habitats, including soils^7,8^, air^9^, aquatic environments^10^, plants^11,12^ and sediments^13^. Recent studies have revealed not only the widespread presence of ARGs, but also their extensive diversity reflecting a broad array of resistance mechanisms to different antibiotic classes^14^. In this context, human activities were shown to contribute significantly to the proliferation, diversification, and especially the environmental mobility of these resistance elements^13,15^.

Among the wide range of anthropogenic drivers, agricultural practices could play a key role in shaping the dynamics of AMR in the environment for several reasons. First, the use of antibiotics as growth promoters or for prophylactic and therapeutic purposes in animal husbandry exerts strong selective pressure on the gut microbial communities of farm animals, promoting the emergence and persistence of both ARB and ARGs^16,17^. In addition, the use of trace elements as feed additives, particularly copper and zinc, for their veterinary properties and growth-promoting effects can promote co-selection and amplification of antimicrobial resistance^18,19^. These resistance determinants can be excreted in animal waste and subsequently introduced into the environment through the application of manure or slurry as fertilizer, contaminating soil, water bodies, and crops^20,21^. Similarly, in agroecosystems, the use of antibiotics, contaminated urban wastes, such as sewage sludges and treated irrigation wastewaters can lead to ARG dissemination within the phyllosphere and rhizosphere microbiota^22^. The above practices not only contribute to the enrichment of ARGs in agricultural soils, but also increase the likelihood of horizontal gene transfer (HGT) between environmental, commensal, and potentially pathogenic bacteria^6,23^. In this context, the close interactions between humans, animals, plants and their shared environments in agricultural settings facilitate transmission routes, blurring the boundaries between environmental, animal and human AMR reservoirs^24–26^. This interconnectedness highlights the importance of adopting a One Health approach to better understand and mitigate the role of agriculture in the global AMR crisis^27^. In this context, a wide array of methodological approaches has emerged to quantify and characterize ARGs across diverse compartments, including culture-based assays, molecular techniques (e.g., MALDI-TOF, qPCR, DNA sequencing, metagenomics), and integrative modeling tools^28^.

In this study, we aimed to conduct a preliminary assessment of the dissemination of antibiotic resistance genes (ARGs) within the environment, focusing on different compartments of tropical agroecosystems across Réunion Island. This tropical island offers a unique context for conducting this study: although geographically isolated, it maintains strong connections with other island territories in the southwestern Indian Ocean, positioned between Southern Africa and the Indian subcontinent, as well as with Europe through its membership of this political community. Réunion benefits from a healthcare system equivalent to that of mainland France and is subject to European agricultural regulations. In addition, several local farming practices—such as the use of livestock manure, the reuse of treated wastewater for irrigation, and the spatial proximity between animal husbandry, crop production and human habitats—could contribute to the circulation of resistance genes in the environment. Previous studies have reported the presence of antibiotic-resistant bacteria in hospitalized patients, pig/poultry farming systems as well as in wastewaters and coastal waters^24,29–32^. However, the occurrence of ARGs in environmental and agricultural matrices remains largely unexplored on the island, as does the potential role of these environments in the dissemination of such genes.

## Material & Methods

### Study sites & sampling

Réunion is a volcanic island constituting a French overseas territory with an area of 2,512 km^2^ and a population of approximately 900,000 inhabitants, located in the south-western Indian Ocean, 679 km from Madagascar and 170 km from Mauritius. Reunion Island enjoys a tropical climate with hot, rainy summers (December to April) and cooler, drier winters (May to November). The topography of Réunion island is highly rugged, with distinct altitudinal zonation: densely urbanized coastal areas give way to mid-altitude agricultural areas (150-600 m) where sugar cane and livestock farming predominate, before transitioning to a national park encompassing much of the island’s interior.

Sixteen samples were collected in June 2022 from various matrices and agroecosystems across the island, as summarized in **Figure 1**. Site A (Soere Pro Reunion) is a 1-ha long-term field trial established in 2014 and located in the northern region (Sainte-Marie), investigating the partial substitution of imported synthetic fertilizers by locally-produced organic fertilizers in a sugarcane agroecosystem^33^. Site B is a watershed located in the southwest (Petite-Île), covering approximately 850 ha and comprising around one hundred smallholder farms practicing livestock farming, crop production (grassland, market-gardening crops, and sugarcane along an approximative decreasing altitudinal gradients) or both. Site C, located in the east (Salazie), is a semi-intensive farm in an area characterized by intensive livestock activity, where poultry and pigs are bred. Animal manure is applied to fertilize one hectare of land dedicated to vegetable production. Although we did not have access to a detailed list of the types and quantities of pharmaceuticals used on the different sites and farms sampled, surveys conducted during sampling confirmed that the livestock producing the manure received occasional antibiotic treatments.

**Figure 1.**
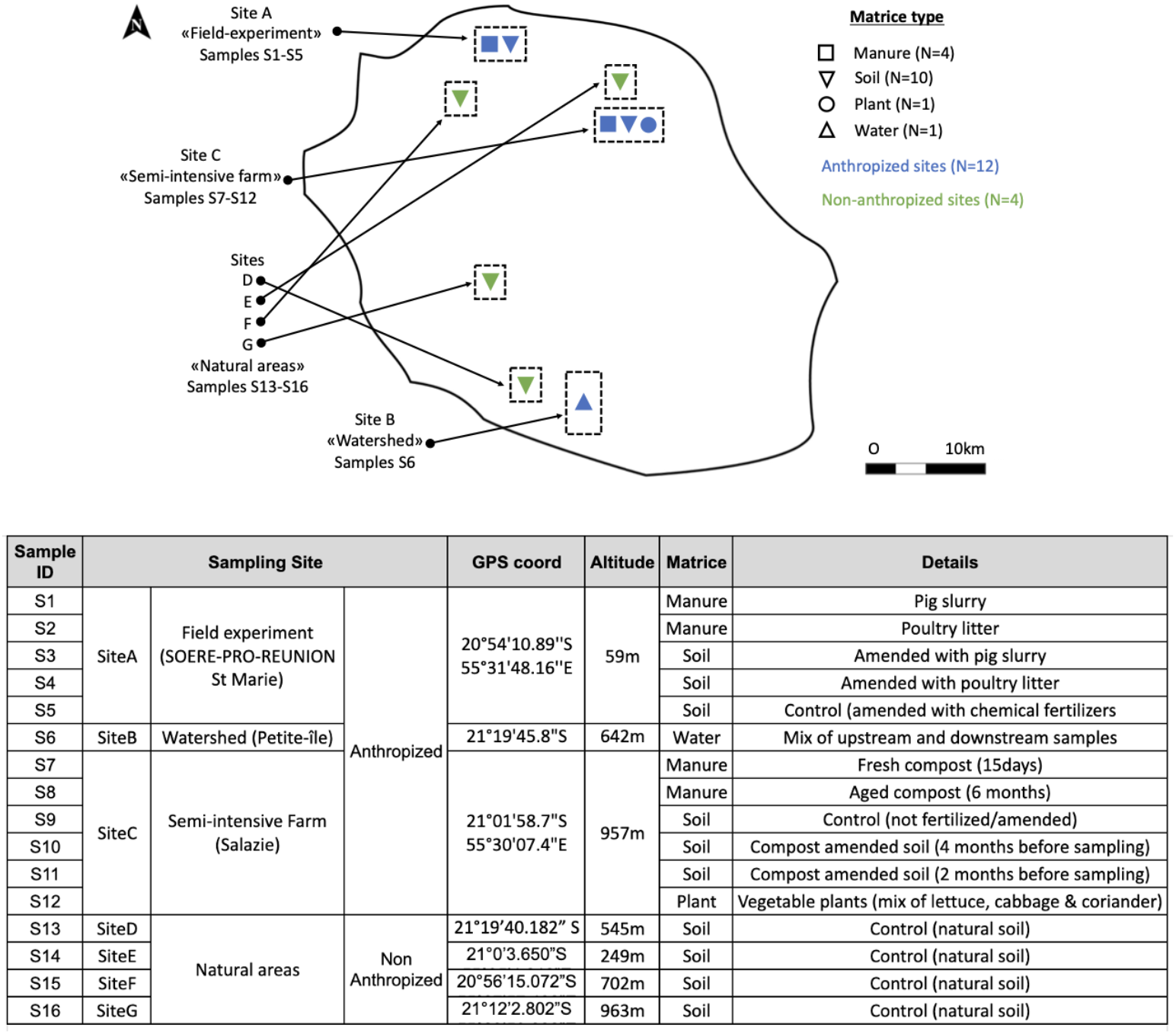
Description of sampling sites and environmental/agricultural samples collected across Réunion Island.

At Site A, five samples were collected, including pig slurry, poultry litter, and soils amended with each of these organic fertilizers, along with a control soil fertilized only with synthetic fertilizers. Site B provided one water sample. At Site C, six samples were obtained: fresh manure (15 days aged) and a 6-month composted manure from a mixture of pig slurry, pig manure and poultry litter, soils either unfertilized (control) or amended with compost four or two months prior to sampling, and vegetable plants grown in plots fertilized with compost. These samples (S1–S12) originated from agroecosystems and are consequently “anthropized”. In contrast, four additional control soil samples (S13–S16) were collected from natural areas (Sites D, E, F, and G) on the edge of or within the La Réunion natural park that had not been subject to any recent agricultural or human influence. These samples have been consequently considered as non anthropized.

In total, four distinct matrices were sampled: manure (S1, S2, S7, S8), soils (S3-S5, S9-S11, S13-S16), vegetable plants (S12) and water (S6). Soils were categorized as follows: *i)* “anthropized” and amended with animal manure (S3, S4, S10, S11), *ii)* “anthropized” and non-amended with animal manure (S5 & S9; i.e. anthropized controls) or “non-anthropized” (S13-S16; i.e. control natural soils).

Sampling was conducted consistently according to the following protocol. For soil sampling, six subsamples were collected from the surface layer (0–10 cm) at various points within each plot using a hand auger, then pooled and sieved through a 4 mm mesh. Manure samples were similarly composed of six subsamples taken from different locations of the manure source and mixed thoroughly. Water samples were collected throughout week 20 of 2022 from two automated stations positioned upstream and downstream of the watershed. Each station collected stream water subsamples every 1 hour 15 minutes, which were pooled to generate the weekly composite samples. A total volume of 3 liters from both stations was pooled and filtered through a 0.2 μm mixed cellulose ester membrane filter (GE Healthcare Life Sciences) under a laminar flow hood. For plant samples, aerial green tissues were collected from fresh lettuce, coriander, and cabbage leaves, which were then chopped and homogenized. All materials were handled with sterile gloves, and equipment was disinfected with 99% isopropyl alcohol between each sample. Samples were transported in ice in a styrofoam cooler and stored at −20 °C in the laboratory until molecular and chemical analyses were performed.

### Antibiotic residues and trace elements screening

A total of six trace elements and 52 antibiotic residues were quantified in all samples using standardized analytical chemistry techniques, with detailed analytical procedures described in **supplementary methods**.

### DNA extraction & gene analysis with high-throughput qPCR arrays

DNA was extracted from the manure and soil samples using the Qiagen DNeasy PowerSoilProKit, using 250mg and 50mg of material, respectively. Qiagen PlantProKit and PowerWaterkits were used to extract DNA from 100mg of plants and water filters, respectively. All extractions were performed according to the manufacturer’s protocols and were stored at −20 °C before qPCR analysis. A total of 384 primer sets (see **File S1**) were used to monitor the presence of 332 ARGs conferring resistance to 10 major classes of antibiotics: aminoglycosides (AMI), β-lactams (BL), multidrug resistances (MDR), macrolide-, lincosamide- and streptogramins B (MLSB), phenicol (PHE), quinolone (QUI), sulfonamides (SUL), tetracycline (TET), trimethoprim (TRI) & vancomycin (VAN), 18 mobile genetic elements (MGE) and 4 integrons (INT) associated with AMR. In addition, 10 genes associated with microbial taxonomic units (TAX) and 3 16S rRNA control genes were also included. Out of the 332 ARGs analyzed, 60 genes were classified as “high risk for human health” based on a previous study^34^ (**File S1**).

Following normalization, gene detection and quantification were carried out using the SmartChip™ Real-Time PCR system (Takara Bio, CA, USA) under the cycling conditions described previously^21^. Briefly, the qPCR conditions included initial enzyme activation at 95°C for 10 min, 40 cycles of denaturation at 95°C for 30s and then annealing at 60°C for 30s for amplification. Each DNA sample was analyzed in three qPCR reactions (i.e. technical replicates). Melting curve analysis was performed for each primer set from all samples. Amplicons with unspecific melting curves and multiple peaks based on the slope of melting profiles were considered to be false positive data and were therefore discarded from the analysis. Melting curve analysis was processed using the SmartChip™ qPCR software.

### Data and statistical analysis

Data analysis including visualization in figures and statistical analysis were performed using R (4.4.2)^35^, the tidyverse ecosystem^36^, ggplot2^37^ package and RStudio (2024.12.0)^38^. A cycle threshold (*Ct*) of 27 was set as the limit for gene detection^21^ and a mean *Ct* value was calculated when a gene was detected in at least two technical replicates. For each replicate sample, the relative gene abundance (R) in proportion to the 16S rRNA gene was computed using the comparative CT method from the following equation^39^:

*R* = 2^−Δ*Ct*^, with Δ*Ct* = *Ct*_(*ARG*)_ − *Ct*_(16*S rRNA*)_ and *Ct* values being averaged over technical replicates.

For paired samples including both a treatment and a control, fold change in ARG abundance (FC) was calculated using the following equation^39^:

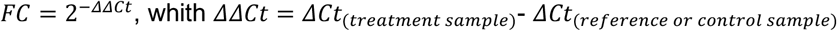

We applied non-metric multidimensional scaling (NMDS; using the `metaMDS` function from the *vegan* package^40^) based on Bray–Curtis dissimilarities to visualize complex multidimensional data, in this case the abundance patterns of antimicrobial resistance genes (ARGs), in a reduced number of dimensions. Finally, we performed four independent linear regression analyses to evaluate the relationships between trace element residues and ARG numbers or relative abundances.

## Results & discussion

### Taxonomic genes

Out of the 10 target taxonomic genes, *Campylobacter* was the only genus absent from all samples (**Figure S1**), suggesting that it may not be a prevalent bacterium in the environments studied. In contrast, members of the phyla *Bacteroidetes* and *Firmicutes* were consistently detected in all samples, highlighting their widespread distribution across both animal-associated and environmental habitats, including soils, plants, and water. Notably, *Streptomyces* were detected exclusively in natural soils, indicating a possible link to undisturbed or native microbial communities, while *Enterococci* were detected exclusively in manure samples, indicating a preferential association with the animal-associated microbiota. The remaining target taxa including *Acinetobacter baumannii, Klebsiella pneumoniae, Pseudomonas aeruginosa*, and *Staphylococci* were detected sporadically, with no obvious link to specific sample types. This variable distribution may reflect differences in environmental persistence, niche adaptation, or recent contamination events^41^.

### Anthropized vs non-anthropized signature of ARGs in control soil samples

When examining only the control samples, we observed no significant difference in both average number and relative abundance of ARGs detected between control anthropized (i.e. unamended) (S5 & S9) and non-anthropized (i.e. natural) (S13–S16) soil samples (**Figure S2**). This result contrasts with previous studies that reported higher numbers and abundance of ARGs in anthropized environments compared to pristine environments^42^. Such a discrepancy could stem from the high diversity and specificity of tropical soils^43,44^ or from the possibility that our supposedly non-anthropized sites are not as isolated as they seem due to their proximity to human activities or historical activities in these regions (the natural park was created in 2007) about which we have no information. As a consequence, all control samples, both anthropized and non-anthropized, were merged for subsequent analyses.

### Number, diversity, and abundance of detected ARGs

On average, 190 ARGs were detected in all samples, but a total of 310 out of 371 ARG qPCR assays were positive in one or more samples (**File S1**). Of the positive assays, 296 targeted ARGs and 14 MGEs. The highest number of ARGs (N=256) was found in sample S3 (soil amended with pig slurry), followed by sample S8 (N=243, aged compost) and sample S7 (N=230, fresh compost). The lowest number of ARGs was found in sample S6 (N=89, water) (**Figure 2A**). On average per matrix type, the highest number of genes was found within the soils amended with manure (N=220, sd=24) and the manure itself (N=219, sd=24), then followed by control soils (N=175, sd=6), vegetable plants (N=143) and water (N=89) (**Figure 2B**). Amongst the 310 ARGs detected, 82 (26%) were found in all the five matrices; 36 (11%) were unique to manure; 17 (5%) were unique to amended soils; 4 (1%) were unique to control soils; and 1 was unique to vegetable plants (**Figure 2C** and **Table S1**). ARGs abundance was quantified relative to the 16S rRNA gene. Only two genes displayed values greater than one, indicating higher abundance than the reference gene. These were “trbC” and “Tn5403”, two genes from the MGE family identified within sample S3 (soil amended with pig slurry). On average, the highest and lowest abundances were observed in sample S3 (soil amended with pig slurry; 0.038) and S6 (water; 0.0008), respectively (**Figure 3A**). On average by matrix type, the highest abundance was found within amended soils (R=0.021, sd=0.01), followed by manure (R=0.010, sd=0.001), control soils (R=0.004, sd=0.0005), vegetable plants (R=0.003) and water (R=0.0008) (**Figure 3B**). Surprisingly, our findings differ from several previous studies that used similar qPCR-based methodologies and reported higher numbers and relative abundance of antibiotic resistance genes (ARGs) in organic manure than in the soils to which it was applied^21,45,46^. This discrepancy could be partly explained by the characteristics of the tropical soils in our study area, which may be richer in microorganisms than the subtropical and temperate soils examined in earlier research, potentially influencing ARG persistence and detection^43,44^. Results of a recent study conducted at the same site showed that soil microbial biomass increased significantly, by an average of 35–45%, in organically fertilized plots, up to 3 to 5 years after the initial application of organic amendments^47^.

**Figure 2.**
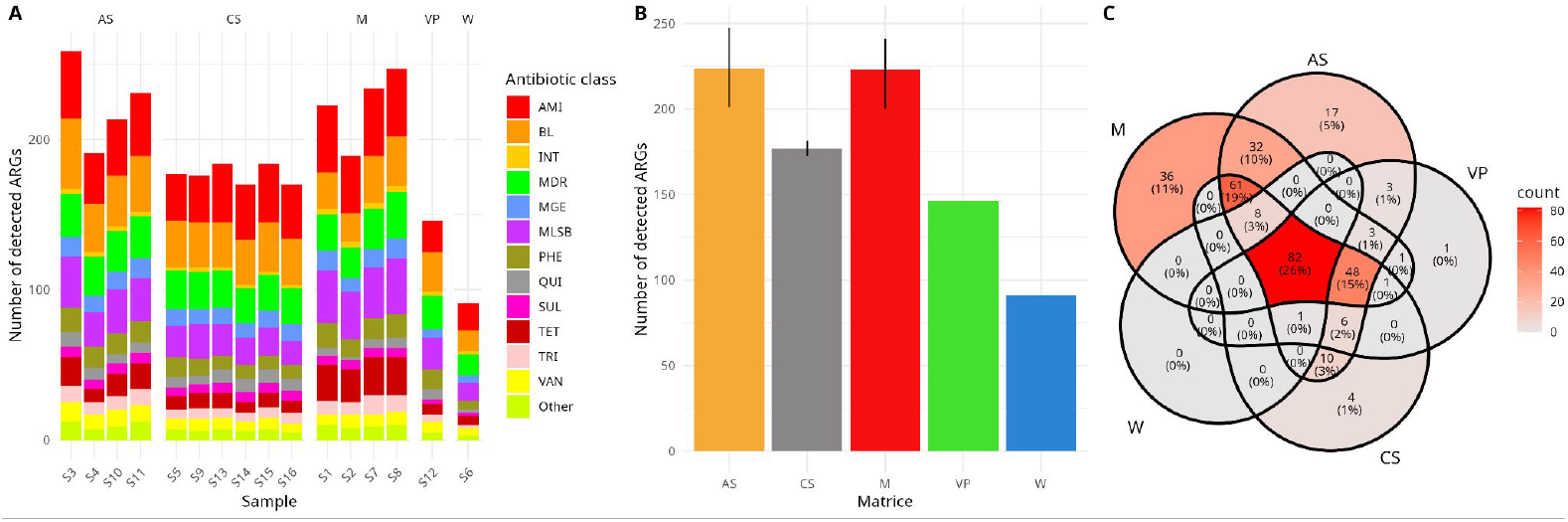
Number and distribution of detected ARG amongst samples (A) and matrices (B). Error bars indicate 95% confidence intervals estimated by nonparametric bootstrap resampling. Venn diagram showing the distribution of antibiotic resistance genes (ARGs) detected across the different matrices, with overlaps indicating genes shared between matrices (C). AS: Anthropized soils, CS: Control soils, M: Manure, VP: Vegetable plants & W: Water.

**Figure 3.**
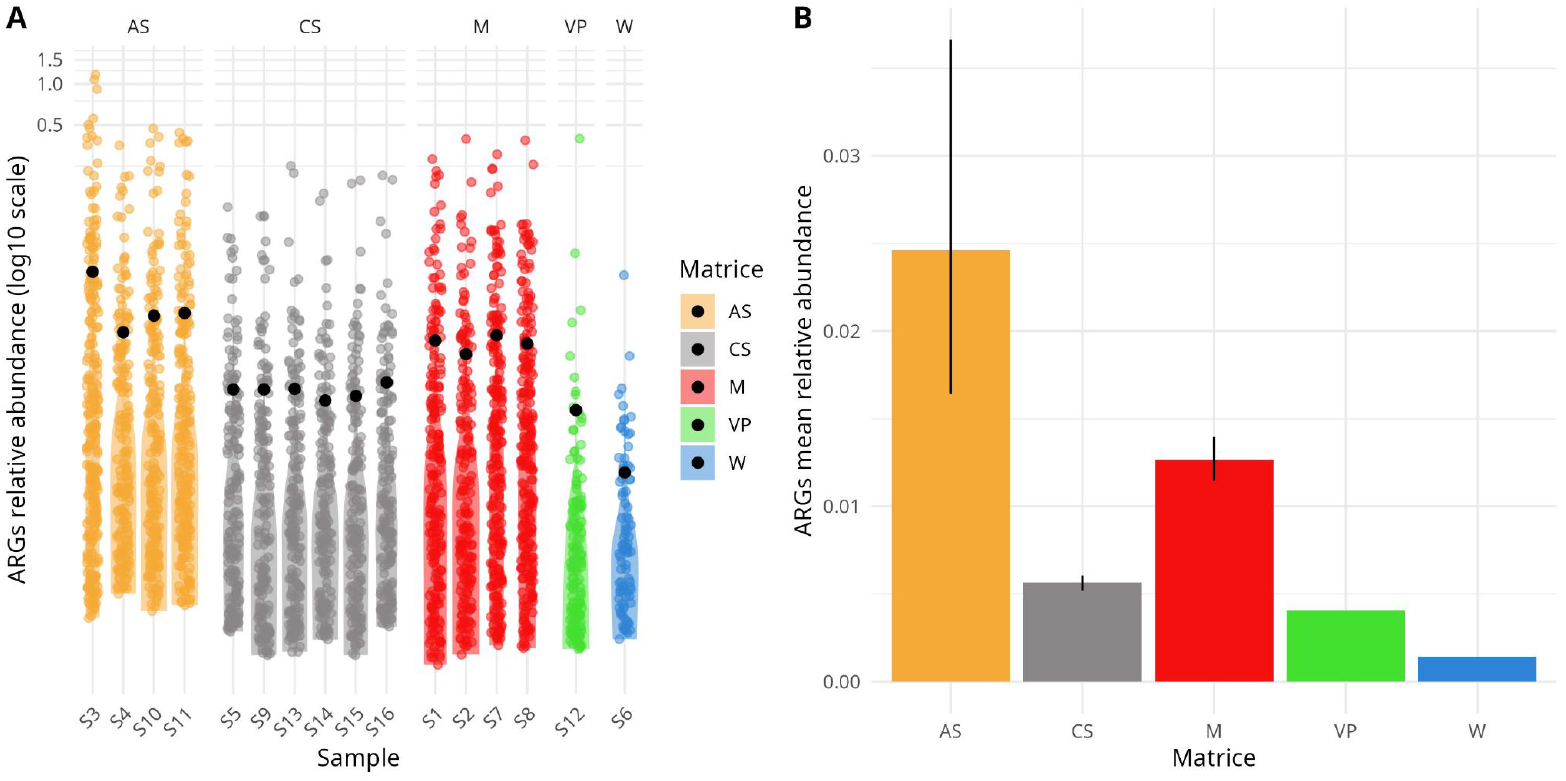
Relative abundance of detected ARG amongst samples (A) and matrices (B). Error bars indicate 95% confidence intervals estimated by nonparametric bootstrap resampling. AS: Anthropized soils, CS: Control soils, M: Manure, VP: Vegetable plants & W: Water.

Globally, ARGs associated with all antibiotic families were detected in all sampled matrices (**Figure 4 & File S1**). ARGs encoding resistance to aminoglycosides, MLSB, quinolone, sulfonamides, tetracycline and trimethoprim were most common in manure and fertilized soils. On the contrary, ARGs conferring resistance to β-lactams were mostly detected in fertilized soils but not in manure. Surprisingly, the catQ gene (associated with resistance to phenicol) was detected in high quantities in plants. The few ARGs found in water were detected in very low quantities compared to other matrices. Unexpectedly, 86% (52/60) of ARGs posing a high risk to human health^34^ were detected in at least one sample and 23% of them (14/60) were found in all samples (**Figure 4 & File S1**). A total of 16 and 22 high-risk antibiotic resistance genes (ARGs) were detected in water and raw vegetable samples, respectively, highlighting a potential public health concern due to the risk of human exposure through contaminated irrigation water, untreated drinking water or the consumption of fresh vegetables. Such a trend has already been observed in other agricultural settings where recycled water or manure-based fertilizers are used, reinforcing concerns about the environmental transmission of antimicrobial resistance throughout the food chain^11,12,48^.

**Figure 4.**
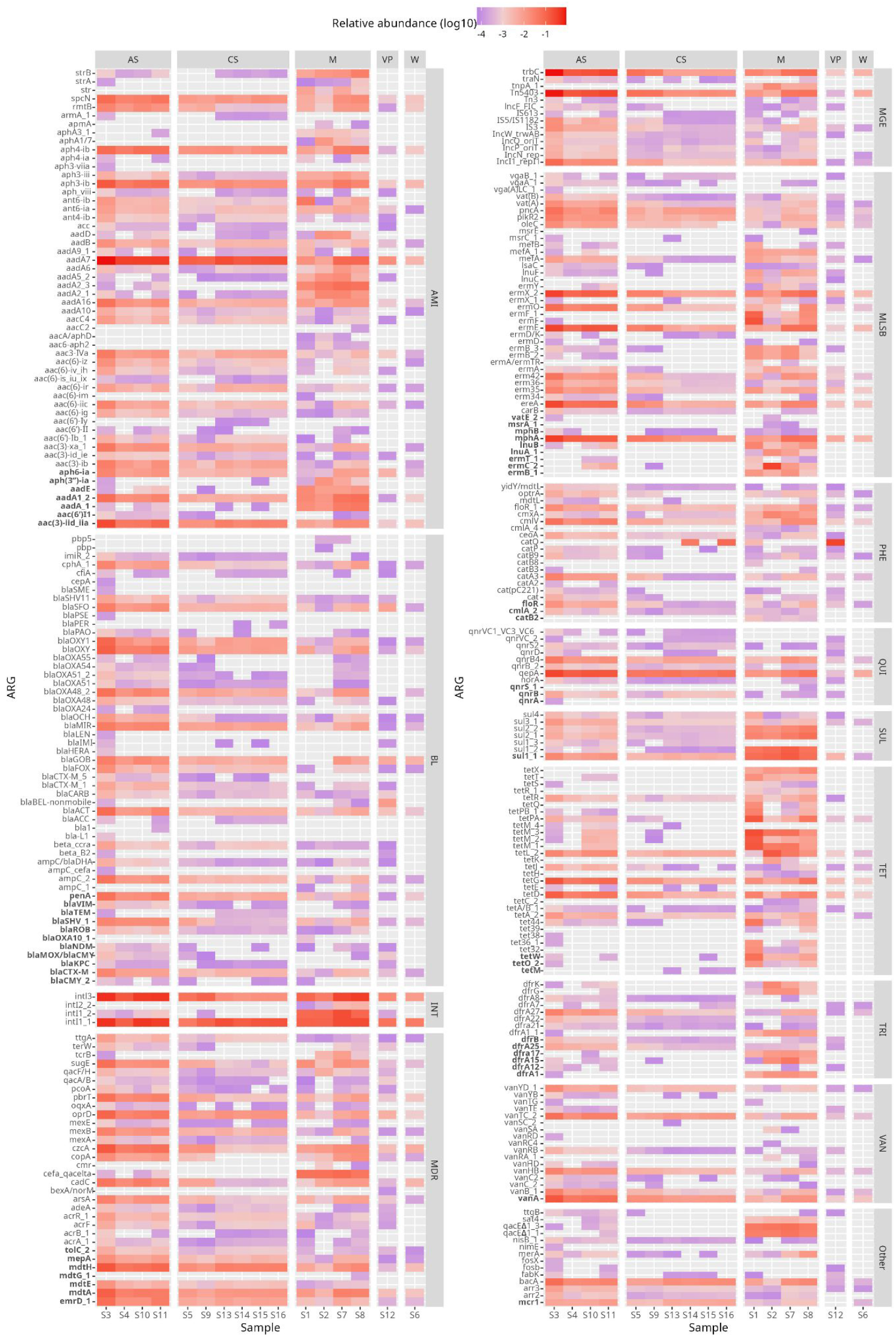
Heat map of relative abundance for 310 ARGs detected in at least one sample. Genes are grouped according to the antibiotic group which the genes confer resistance. ARGs considered “high risk to human health” are highlighted in bold. AS: Anthropized soils, CS: Control soils, M: Manure, VP: Vegetable plants & W: Water.

### Dissemination of ARGs

A total of 65 ARGs were detected in both manure and the amended soils (S1-S3, S2-S4 & S8-S10/S11), while being absent in anthropized control soils (i.e. unamended soil S5&S9), suggesting potential transmission from manure to soil (see **Table S2**). These were mainly ARGs from the MLSB (N=20) and tetracycline (N=12) families. Similarly, we identified 4 potential ARGs from the MLSB and BL families that could have been disseminated from manure to vegetable plants. Interestingly, 10% (7/69) of the ARGs that showed dissemination patterns were on the list of genes posing a high risk to human health list (**Table S2**).

### Changes in ARG abundances due to fertilization, composting and time

On average, ARGs relative abundance was 4.9 times higher in amended soils than in control soils, highlighting the effect of organic amendments on the enrichment and dissemination of antibiotic resistance genes within soil microbial communities in Reunion island. Fold-changes were calculated for the paired-samples described hereafter (**Figure S3**). During composting (measured by comparing S7 vs S8), ARG average relative abundance decreased by 1.6-fold, a trend that has also been reported in previous studies^48–52^. The highest decrease rates (up to 12-fold) were observed for the ermB_3, tcrB and aadF genes of the MLSB, MDR and AMI families, respectively. Although an overall decrease was observed on average, the abundance of certain genes increased during composting. The most pronounced increases—up to 12-fold and 11-fold—were recorded for the tetQ and tetPA genes, both belonging to the tetracycline resistance family. The effect of manure amendment on ARGs abundance in soils was further assessed by comparing amended soils and control soils from the same sites (S3 vs S5, S4 vs S5 & S11 vs S9). On average, the relative abundance of ARGs in amended soils was increased by 7-fold, in accordance with previous studies reporting 2 to 100-fold increases in ARG levels following manure application^21,46,48,52^. The highest increase (up to 75-fold) was observed for the ermC_2 gene, followed by the tetM_2&tetM_3 genes (36-fold) and aadE (25-fold) genes. Finally, when comparing soil samples taken 4 months after manure amendment (S11) and 2 months after (S10), we observed an average decrease in the relative abundance of ARGs over time by 1.5-fold, as previously described^21,53^. Only one ARG (tetPB_1) from the tetracycline family displayed an increase in relative abundance over time of more than 2-fold.

### ARG structuration within contrasted agroecosystems

The two-dimensional NMDS clustering of ARG abundances yielded a stress value of 0.07, indicating a good ordination with no real risk of drawing false inferences^54^. The structuration of ARG abundance showed no association with sampling site, but rather with the nature of the agricultural sample, as similar sample types cluster independently of their geographic origin (**Figure 5**). This pattern may be explained by the fact that the physicochemical properties and microbial communities associated with specific sample types (e.g., soil, manure, plant, water) exert a stronger influence on the composition of ARGs than spatial location. Interestingly, control and amended soils showed a clear genetic structure. Amongst the amended soils, sample S3 (soil amended with pig manure on sampling site A) appears distinct, suggesting slight differences in its ARG abundance profile compared to samples S4, S10 and S11. Both water (S6) and plant (S12) samples, although unique, also showed distinct profiles from the other matrices. Manure samples exhibited some substructure, with fresh (S7) and composted (S8) pig manure clustering together, while chicken (S2) and pig (S1) manure from sampling site A remained distinct.

**Figure 5.**
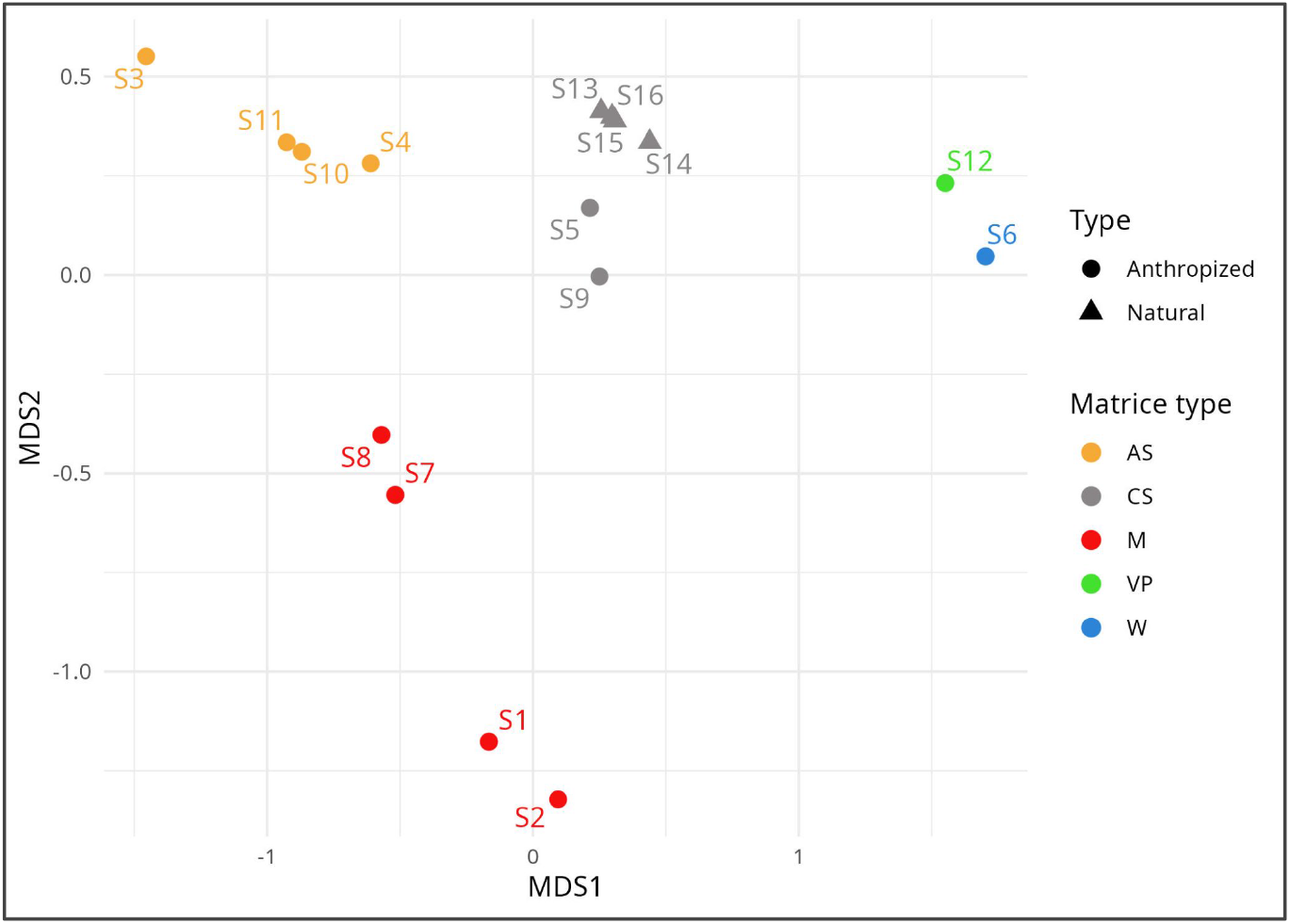
Clustering of different sample matrices in nonmetric multidimensional scaling (NMDS) plot.

### Antibiotic and trace elements residues

Amongst the 52 antibiotic residues tested, only 5 compounds, belonging to the β-lactam (BL), macrolide-lincosamide-streptogramin B (MLSB), quinolone (QUI), and sulfonamide (SUL) families, were detected in 5 of the 16 analyzed samples (**Table 1**). These detections were obtained in S1 (pig slurry, site A), S2 (poultry litter, site A), S8 (aged compost, site C), S9 (anthropized control soil, site C), and S11 (soil amended with compost two months prior to sampling, site C). No antibiotic residues were detected in any of the water, plant, or natural soil samples. These findings are consistent with the very preliminary results obtained on the site A at the beginning of the field trial, where only two antibiotic residues were recovered in animal manures but not in the amended soils^55^. These findings are even more consistent with previous observations made in China, where manure-amended soils and composts are often identified as the primary reservoirs of antibiotic residues^56^. The presence of sulfamerazine in the control soil sample S9 may be attributed to contamination from a nearby amended field, potentially resulting from agricultural activities or runoff water, as suggested by the detection of the same molecule in the close-by amended soil sample S11. The absence of detectable residues in water and plant compartments may be explained by several non-exclusive mechanisms: rapid biotic or abiotic degradation, strong sorption to soil organic matter and minerals, significant dilution by uncontaminated waters at the watershed scale, high temporal variability in antibiotic concentrations related to episodic inputs, or inherently low uptake and translocation potential of antibiotics in plants. In addition, the lack of detectable residues in natural soils reinforces the hypothesis that point sources such as manure and compost are the dominant contributors to antibiotics environmental contamination.

**Table 1.** Antibiotic traces detected. Values shown in grey designate concentrations expressed in µg/kg, whereas white cells represent measurements below the indicated detection limit. Only samples and antibiotics with at least one detectable value are included.

| Antibiotic | Family | Samples |  |  |  |  |
| --- | --- | --- | --- | --- | --- | --- |
|  |  | S1 | S2 | S8 | S9 | S11 |
| Dicloxacillin | BL | <1,96 | 15,4 | <1,96 | <1,96 | <1,96 |
| Tiamulin | MLSB | 9,45 | <3,3 | 26,4 | <3,3 | <3,3 |
| Oxolinic acid | QUI | 609 | <5,8 | <5,8 | <5,8 | <5,8 |
| Sulfamerazine | SUL | <2,8 | <2,8 | <2,8 | 47,8 | 80,1 |
| Sulfamethizole | SUL | <3,0 | <3,0 | <3,0 | <3,0 | 35,7 |

The quantification of trace elements across the different sample types revealed distinct concentration patterns between metals and matrices (**Figure S4**). The elevated Cu and Zn contents in animal manures, compared to other matrices (i.e. soils, vegetable and water), are attributable to the supplementation of these two elements to livestock feed. Only a small fraction of Cu and Zn is metabolized by the animals, with most being excreted in livestock effluents and subsequently applied to soils, where Cu and Zn generally accumulate in the surface soil layers ^57,58^. In this study, total Cu and Zn concentrations in natural soils did not differ significantly from those in anthropized soils. This reflects the naturally high background concentrations in Réunion Island soils derived from volcanic parent material^59^, which masks the contribution of livestock effluent inputs. Consequently, we characterized the DTPA-extractable fractions of Cu and Zn, which were markedly higher in animal manures (Figure S1). In natural soils, the mean DTPA-extractable Zn fraction was significantly lower than in anthropized soils (control and amended). A similar nonsignificant trend was observed for Cu. These findings suggest that fertilization and amendment practices have increased the amount of Cu and Zn in anthropized soils relative to natural soils.

### Association between antibiotic residues, trace elements and ARGs

Because of the small number of samples in which antibiotic residues were detected (**Table 1**), we were unable to establish a statistical correlation between antibiotic concentration and ARG abundance. Therefore, for each antibiotic identified, we examined the distribution of all known associated ARGs. Across all samples, we observed no consistent association between the presence or amount of a given antibiotic and the abundance of the corresponding resistance genes (**Figure S5**). For the trace elements, no significant correlation was observed between the combined concentrations of copper and zinc and either the number or abundance of ARGs at a 5% significance threshold (**Figure S6**). This apparent discrepancy could be explained by the fact that many ARGs may originate from a compartment other than the one in which they were ultimately detected, after being introduced by environmental transport, host migration, or horizontal gene transfer. Such lack of correlation has previously been reported in multiple studies, suggesting that ARG persistence and dissemination can occur independently of local selective pressure^46,60,61^. This underscores the complexity of ARG dynamics and the importance of considering both historical and spatial dimensions of resistance gene propagation within a One Health context.

In conclusion, this study provides the first exploratory assessment of antibiotic resistance genes (ARGs) and antibiotic residues across diverse agricultural matrices on Réunion Island. Our results highlight that organic fertilizers such as manure and compost contribute significantly to the enrichment and dissemination of ARGs in soils, with notable detection even in water and vegetable samples, raising potential public health concerns. Interestingly, the structuration of ARG profiles appears to be more strongly influenced by sample type than by geographic origin, suggesting that specific physicochemical properties and microbial communities associated exert a stronger influence on the composition of ARGs than local spatial location. Moreover, we observed no significant association between the detection of trace elements or antibiotic residues and the presence or abundance of associated ARGs, suggesting that the persistence and dissemination of resistance genes in the environment may occur independently of current selective pressure, potentially reflecting past contamination events or horizontal gene transfer dynamics. These results are original, but this exploratory study has some limitations, including the limited number of sampling sites, the inability to link the detected ARGs to their specific bacterial hosts and the lack of direct information on local historical and contemporary antibiotic usage patterns. Future work should expand the spatial and temporal scope of sampling, incorporate either bacterial culture or metagenomic approaches to better characterize the microbial hosts of ARGs, and further investigate the potential transmission pathways between environmental, animal, and human compartments. Such efforts will be essential to inform One Health-based strategies aimed at mitigating AMR in tropical agricultural systems.

## Data availability

The raw qPCR output data, along with the R code used for data analysis and figure generation, are available on GitLabat at the following link: https://gitlab.cirad.fr/pvbmt/2026_AMR_RUN

## Acknowledgments

We thank the various members of producer cooperatives, farmers, and veterinarians who responded to our surveys on antibiotic use in Réunion. We are grateful to the Recyclage et risque group (CIRAD) for the monitoring of the site A over years and more particularly to G. Moussard and C. Detaille for their help in recovering and preparing samples. We extend our thanks to Windi Muziasari and the entire Resistomap team for their assistance with qPCR data production and interpretation. We are grateful to G. Miltgen and D. Wilkinson for valuable discussions regarding the choice of ARGs and their circulation in both human and animals compartments, respectively.

