## Supplementary tables & figures for "Distribution of antibiotic resistance genes across contrasted tropical agroecosystems in Réunion Island"

**Supplementary methods**

Searching for antibiotic residues Antibiotic residue analysis was outsourced to the Resistomap analytical platform for the following 52 residues encompassing major antibiotic classes: β-Lactams (Amoxicillin, Ampicillin, Cloxacillin, Dicloxacillin, Oxacillin, Penicillin, Cefaclor, Cefadroxil, Cefalexin, Cefazolin), Macrolides (Azithromycin, Clarithromycin, Erythromycin, Tylosin), Lincosamides (Lincomycin), Pleuromutilins (Tiamulin), Phenicols (Chloramphenicol, Florfenicol, Thiamphenicol), Quinolones & Fluoroquinolones (Ciprofloxacin, Danofloxacin, Difloxacin, Enrofloxacin, Marbofloxacin, Norfloxacin, Ofloxacin, Sarafloxacin, Flumequine, Oxolinic acid), Tetracyclines (Chlortetracycline, Doxycycline, Oxytetracycline, Tetracycline), Sulfonamides (Sulfachloropyridazine, Sulfaclozine, Sulfadiazine, Sulfadimethoxine, Sulfadimidine, Sulfadoxine, Sulfaguanidine, Sulfamerazine, Sulfamethizole, Sulfamethoxazole, Sulfamethoxypyridazine, Sulfamonomethoxine, Sulfamoxole, Sulfapyridine, Sulfaquinoxaline, Sulfathiazole, Sulfisoxazole) & Diaminopyrimidines (Trimethoprim).

The extraction method for the determination of antibiotic residues (with the exception of β-Lactams) in solid samples was carried out with the procedure detailed by Gago-Ferrero et al. (2015)[1]. The extraction method for the determination of β-Lactams in solid samples was based on the procedure developed by Meklati et al. (2022)[2], with the following minor changes. Three grams of solid samples were weighed and inserted into a 50 mL polypropylene tube, and then a volume of 30 mL of phosphate buffer solution (PBS) was added. This PBS buffer was previously prepared by dissolving 2.176 g of potassium phosphate dibasic anhydrous in 250 mL of water, while adjusting the pH to 8.5 using 0.1 M solution of NaOH. Adjustment of the PBS buffer solution pH to this value is essential for the recovery of penicillins and cephalosporins from the matrix. The samples were shaken onto an horizontal shaker for 15 min. A volume of 1 mL of acetonitrile was added followed by vortex agitation. Centrifugation at 4000 rpm for 10 min was performed and the supernatant was pipetted and transferred in a new tube. The samples as well as the spiked ones were prepared in the same conditions. All samples were subsequently cleaned up by solid phase extraction (SPE, Strata-X cartridges 200 mg/6mL) to remove interferences from the matrix. The cartridges were preconditioned with 10 mL methanol and 5 mL PBS buffer solution (0.05 M; pH 8.5), and then the samples were loaded onto the cartridges. In order to remove interferences, a washing step was performed with 3 mL PBS and 2 mL water. The SPE cartridges were then dried for 5 min under vacuum, and the analytes were eluted using 5 mL acetonitrile. The samples were evaporated to dryness at 40 °C and reconstituted before analysis with 300 µL of 0.1% formic acid: MeOH 0.1% formic acid (95:5, v/v). Finally, the extraction method for the determination of antibiotic residues in water samples was carried out with the procedure detailed by Galani et al. (2021)[3]. The only two changes concerned the initial volume of water used (1 L instead of 0.1 L) and the final volume reconstituted before analysis (0.25 mL instead of 0.5 mL).

After extraction, residue quantification was performed using liquid chromatography coupled with tandem mass spectrometry (LC-MS/MS) as detailed by Gago-Ferrero et al. (2015) and Meklati et al. (2022).. In the analysis of sulfonamides, quantification was performed using spiked samples with the use of Sulfadiazine-d4 as internal standard. For analysis of the other residues, the quantification was performed using spiked samples without the use of internal standards. The recoveries have been checked preparing matrix matched samples.

Searching for trace elements The quantification of trace elements was carried out using two complementary approaches. First, total concentrations for total copper (Cutot), zinc (Zntot), nickel (Nitot), lead (Pbtot), chromium (Crtot) and cadmium (Cdtot) were measured as following. Soil, manure, and vegetable samples were first oven-dried at 40 °C. A 100–150 mg portion of the powdered sample was then calcined at 450 °C for 4 h to eliminate organic matter, followed by complete digestion with a mixture of HF, $HNO_3$, and $HClO_4$, in accordance with ISO 14869-1. Water sample was analyzed after filtration, without prior mineralization. Cu and Zn concentrations were determined using a PerkinElmer NexION 300X ICP-MS, with $^{103}Rh$ as the internal standard. Quality control was ensured through the analysis of certified reference materials, including EnviroMAT EP-L-3 and ES-H-2 (SCP Sciences, Courtaboeuf, France), and SRLS-5 (Ottawa, Canada). Second, in addition to total concentrations, the available fraction of copper (CuDTPA) and zinc (ZnDTPA) was assessed in soil and manure samples only. Samples were extracted with the diethylenetriaminepentaacetic acid (DTPA) method according to Lindsay and Norvell (1978)[4], with. The solution, composed of 5 mM DTPA, 10 mM $CaCl_2$, and 100 mM triethanolamine buffered at pH 7.3, was mixed with soil or manure samples at a soil-to-solution ratio of 1:2 or a manure-to-solution ratio of 1:25. The mixtures were then stirred for 2 h and centrifuged at 1000 g, after which the supernatants were filtered through ashless medium-speed filter paper (8 µm pore size). Solutions were analyzed with a flame atomic absorption spectrophotometer. The DTPA extraction procedure was validated by using an uncertified internal reference soil from Réunion when the extracted Cu and Zn recovery was within the 90–110% range and the coefficient variation on the three replicated internal reference was within 10%.

### Supplementary figures

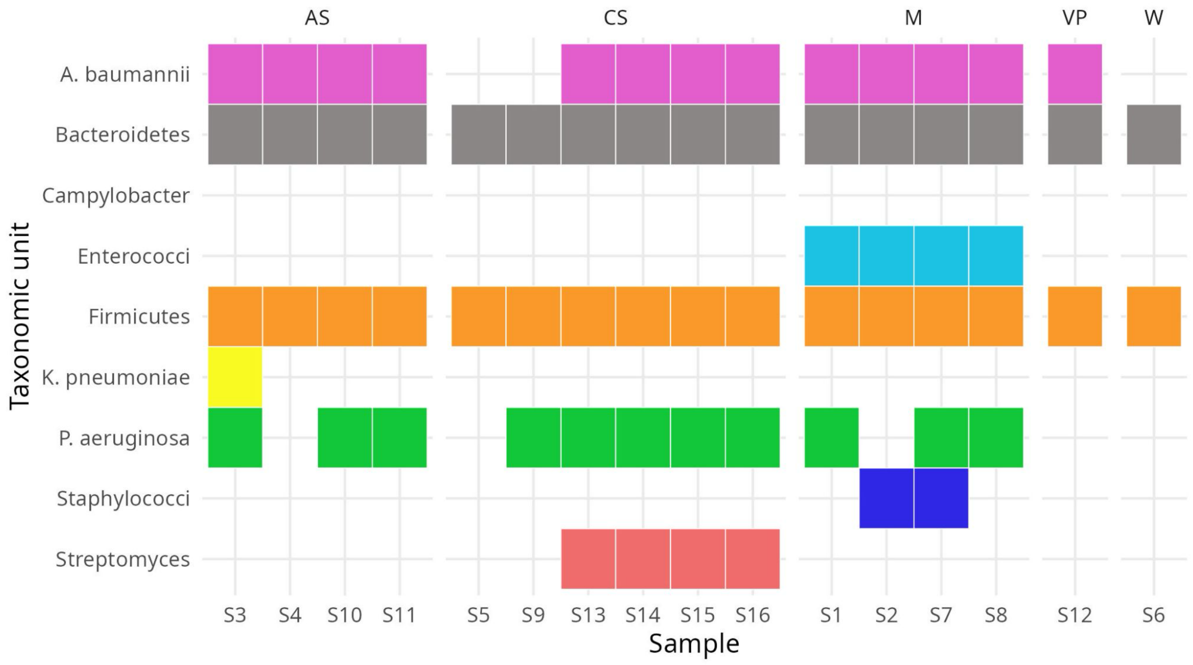

**Figure S1:** Taxonomic composition of each sample

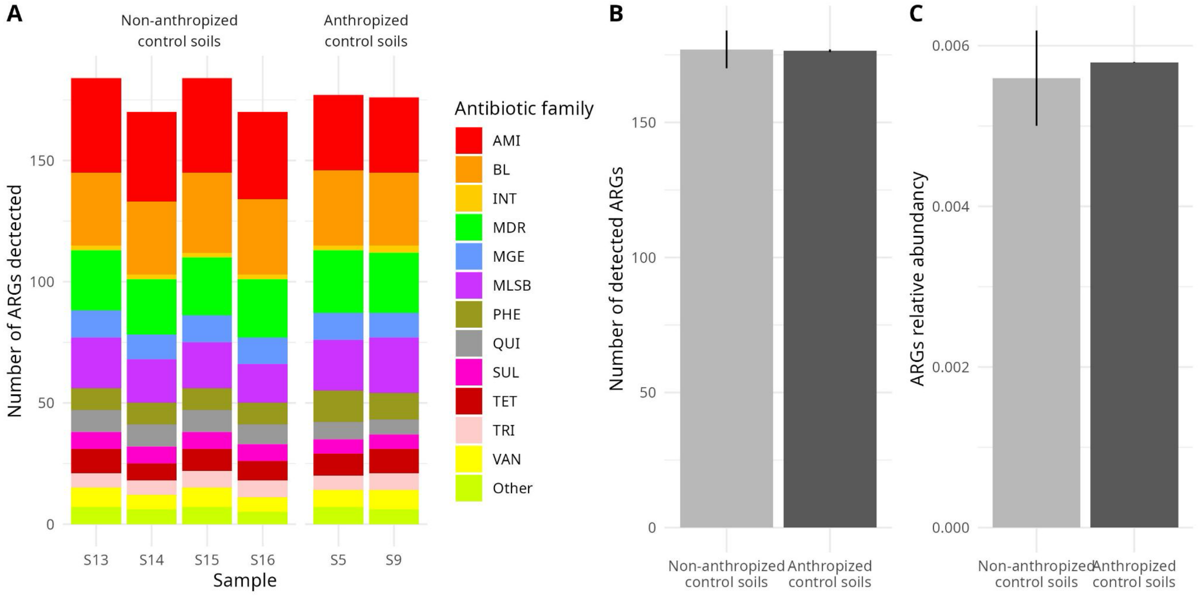

**Figure S2**: ARGs in anthropized vs non-anthropized (natural) control soil samples

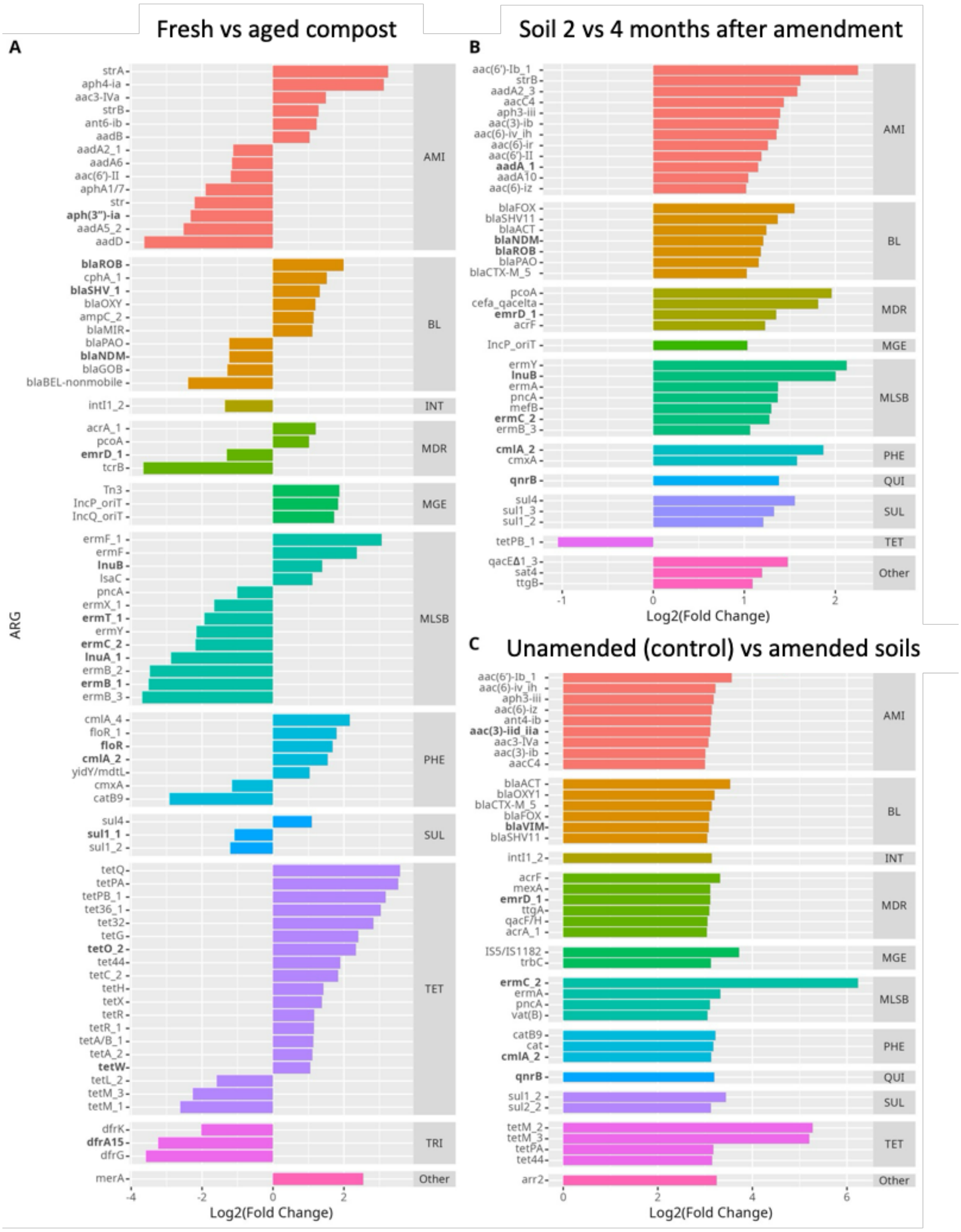

**Figure S3**: Fold change of ARGs between paired samples: fresh to aged compost (A), soil 2 months vs 4 months after amendment (B) and unamended vs amended soils (C). Fold changes are log 2 transformed. Only genes with at least 2-fold changes are shown. ARGs considered "high risk to human health" are highlighted in bold.

| | Nitot | Cutot | Pbtot | Zntot | Crtot | Cdtot | ZnDTPA | CuDTPA |
|---|---|---|---|---|---|---|---|---|
| **S1** | 14,6 | 273,8 | 2 | 1716,7 | 9,1 | 1,01 | 1176,8 | 150,3 |
| **S2** | 7,4 | 112,7 | 0,2 | 647 | 7,2 | 0,36 | 273,2 | 35,2 |
| **S3** | 38,4 | 94,7 | 14,5 | 220,2 | 56,1 | 0,3 | 5,1 | 2,8 |
| **S4** | 32,8 | 83,4 | 10,1 | 190,3 | 47 | 0,22 | 1,8 | 1,7 |
| **S5** | 36,8 | 87,7 | 13,3 | 216,1 | 60 | 0,11 | 5,0 | 3,0 |
| **S6** | 0,4 | 2,1 | 0,5 | 6,4 | 0,8 | 0,03 | NA | NA |
| **S7** | 15,8 | 127,4 | 0,2 | 788,2 | 8,5 | 0,44 | 421,7 | 34,3 |
| **S8** | 19,9 | 125,6 | 0,7 | 715,5 | 17,8 | 0,41 | 414,2 | 27,2 |
| **S9** | 179 | 63,8 | 3 | 130,9 | 211,9 | 0,12 | 2,9 | 1,7 |
| **S10** | 163,7 | 65,3 | 2,4 | 131 | 234,7 | 0,12 | 6,9 | 1,3 |
| **S11** | 155,5 | 63,6 | 2 | 134,7 | 281,6 | 0,1 | 8,4 | 2,1 |
| **S12** | NA | 5 | NA | 50 | NA | NA | NA | NA |
| **S13** | 245,6 | 71,8 | 4,4 | 108,3 | 395,5 | 0,38 | 0,8 | 0,9 |
| **S14** | 189,3 | 37,4 | 5,9 | 97 | 236,2 | 0,01 | 0,6 | 1,5 |
| **S15** | 355,3 | 105,5 | 6,8 | 113,7 | 585,3 | 0,09 | 0,8 | 2,3 |
| **S16** | 48,1 | 28,4 | 10,1 | 199,3 | 67,4 | 0,15 | 1,6 | 1,1 |

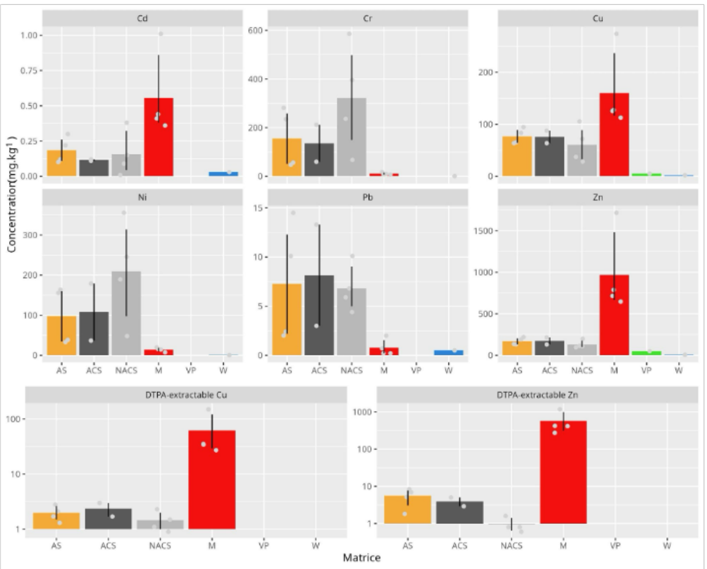

**Figure S4**: Concentrations of total and DTPA-extractable trace elements (Cd, Cr, Cu, DTPA-Cu, DTPA-Zn, Ni, Pb, Zn) across the different samples (left) and matrices (right). (AS: amended soils, ACS: Anthropized control soils, NACS: Non-anthropized (pristine) control soils, M: Manure, VP: Vegetable plants & W: water). Bars show mean ± SD and grey points represent individual measurements.

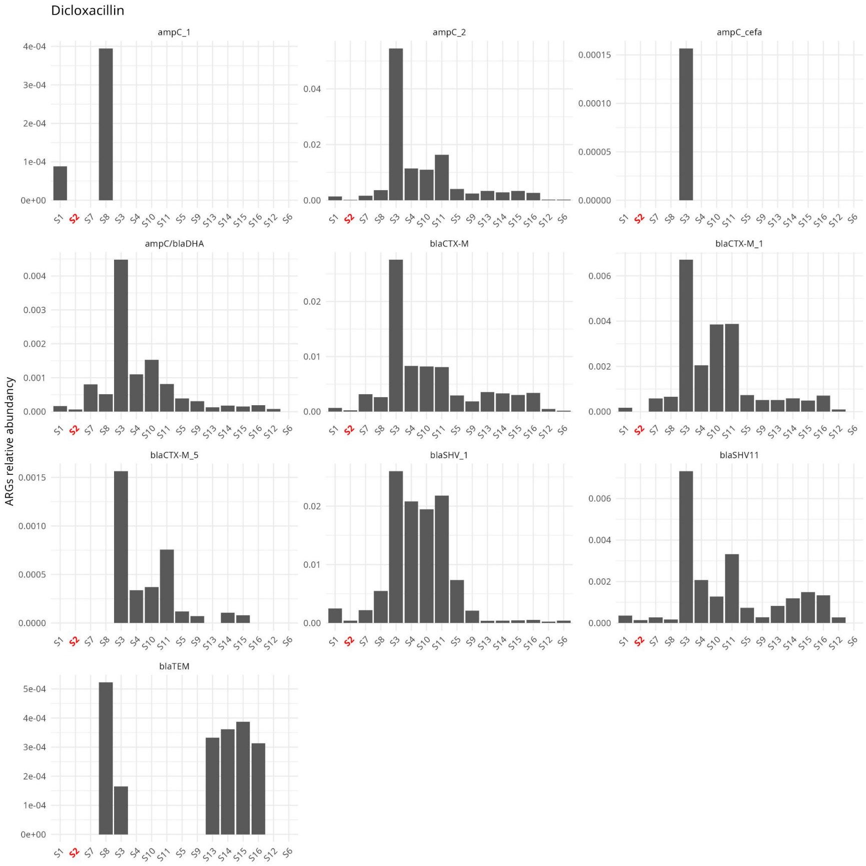

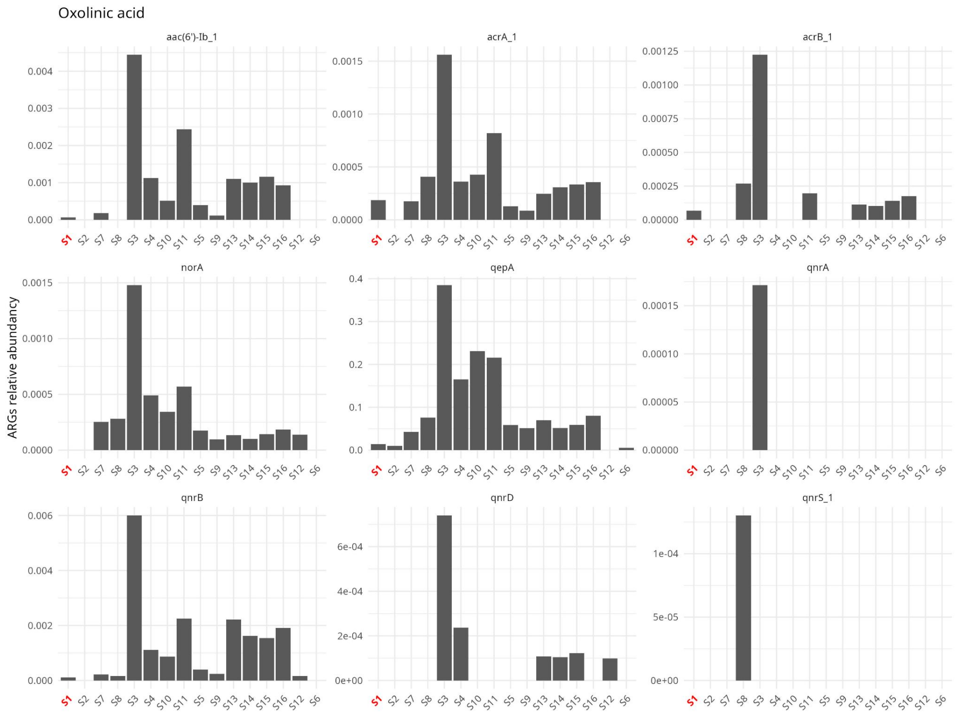
Oxolinic acid
aac(6')-Ib_1
acrA_1
acrB_1
norA
qepA
qnrA
qnrB
qnrD
qnrS_1
ARGs relative abundancy
S1 S2 S7 S8 S3 S4 S10 S11 S5 S9 S13 S14 S15 S16 S12 S6

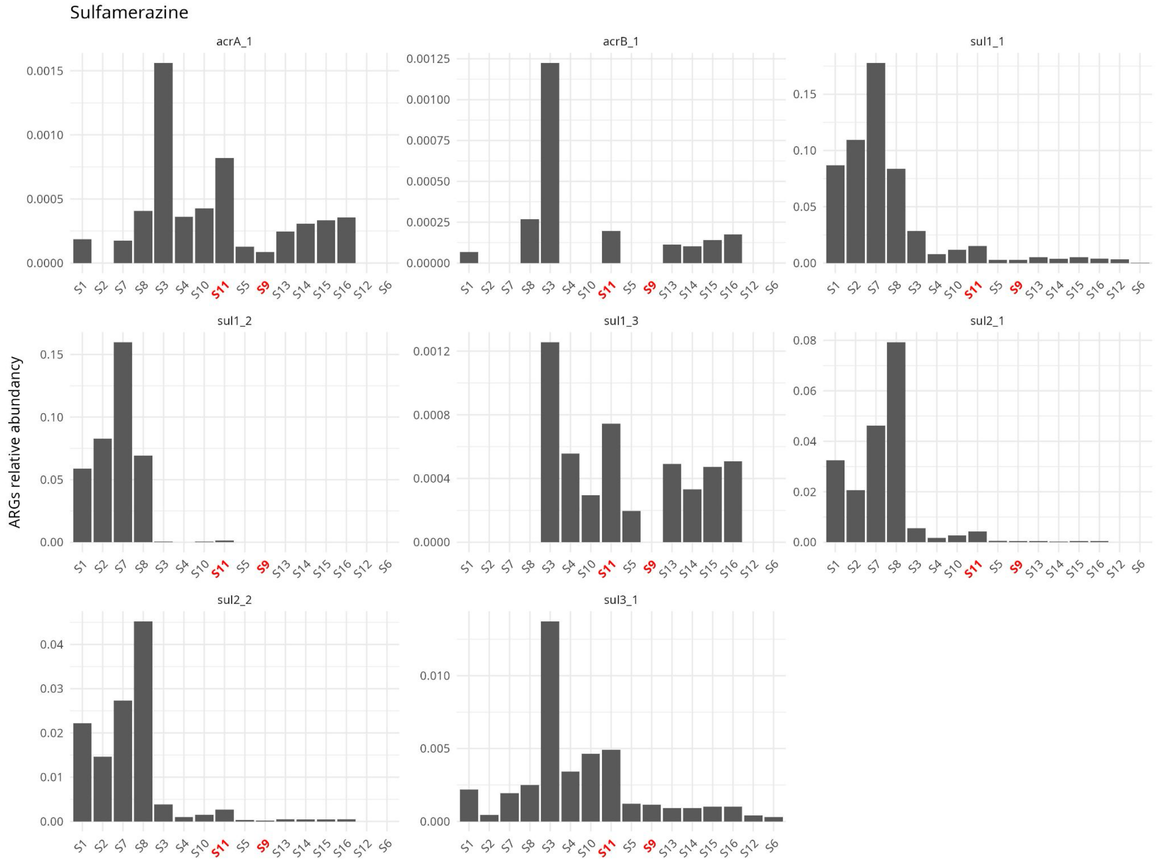
Sulfamerazine
acrA_1
acrB_1
sul1_1
sul1_2
sul1_3
sul2_1
sul2_2
sul3_1
ARGs relative abundancy
S1 S2 S7 S8 S3 S4 S10 S11 S5 S9 S13 S14 S15 S16 S12 S6

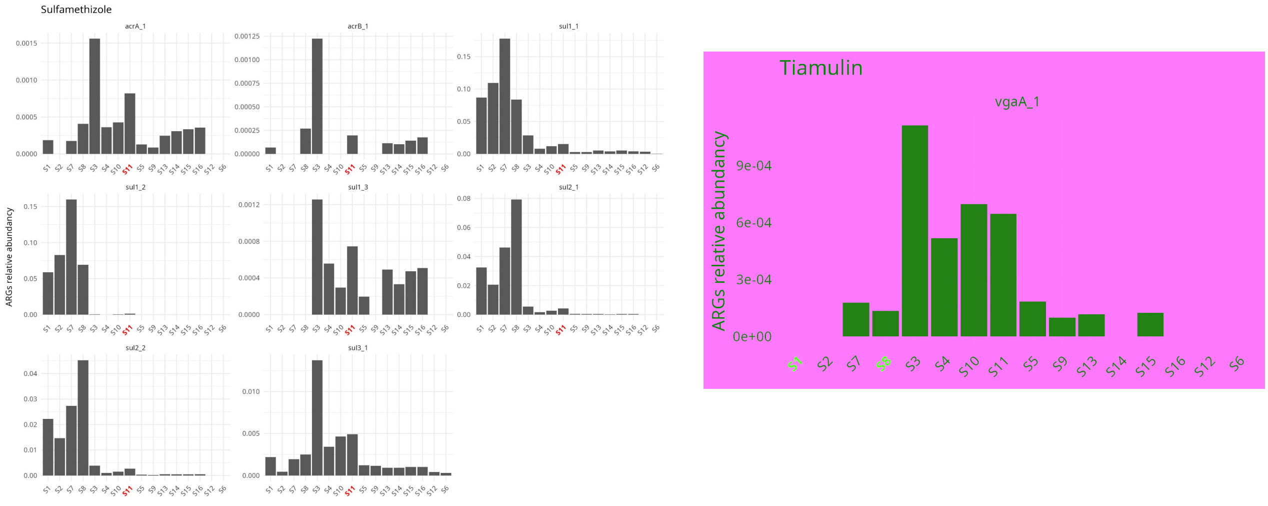

**Figure S5:** Association between antibiotic residues detection and ARG abundance. For each antibiotic, red labels mark the samples where residues were detected.

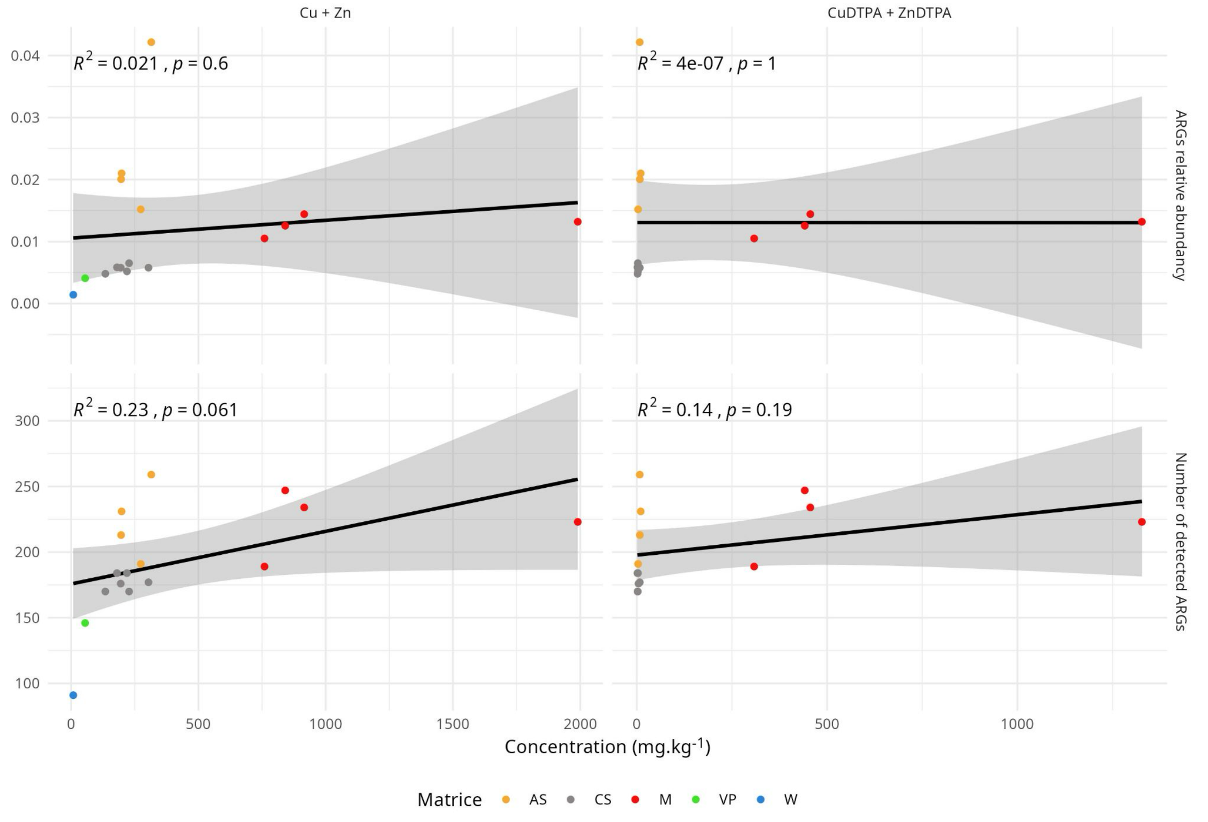

**Figure S6:** Linear regression between Cu/Zn residues and ARG number/abundance. The gray ribbons represent 95% confidence interval of the intercept and slope of each independent regression lines.

### Supplementary tables

**TableS1:** List of genes uniquely detected in specific matrices

| Assay | Gene | Family | Unique to | Assay | Gene | Family | Unique to |
|---|---|---|---|---|---|---|---|
| AY164 | vanRA_1 | VAN | Manure | AY90 | ermA/ermTR | MLSB | Manure |
| AY17 | aphA1/7 | AMI | Manure | AY131 | pbp5 | BL | Manure |
| AY2 | aacC2 | AMI | Manure | AY132 | pbp | BL | Manure |
| AY206 | cmr | MDR | Manure | AY165 | vanSA | VAN | Manure |
| AY212 | mdtG_1 | MDR | Manure | AY176 | vanRC4 | VAN | Manure |
| AY22 | str | AMI | Manure | AY258 | tetK | TET | Manure |
| AY250 | tet32 | TET | Manure | AY66 | msrA_1 | MLSB | Manure |
| AY259 | tetQ | TET | Manure | AY380 | vanSC_2 | VAN | Control soils |
| AY260 | tetH | TET | Manure | AY385 | aac(6')-Iy | AMI | Control soils |
| AY267 | tetX | TET | Manure | AY574 | tetM | TET | Control soils |
| AY268 | tetC_2 | TET | Manure | AY437 | blaPER | BL | Control soils |
| AY294 | intI2_2 | INT | Manure | AY613 | blaOXA24 | BL | Amended soils |
| AY299 | tnpA_1 | MGE | Manure | AY338 | bla1 | BL | Amended soils |
| AY325 | tetR_1 | TET | Manure | AY557 | catA2 | PHE | Amended soils |
| AY406 | aac6-aph2 | AMI | Manure | AY109 | blaPSE | BL | Amended soils |
| AY41 | cmlA_4 | PHE | Manure | AY113 | bla-L1 | BL | Amended soils |
| AY426 | apmA | AMI | Manure | AY115 | cepA | BL | Amended soils |
| AY46 | ermF_1 | MLSB | Manure | AY177 | vanRD | VAN | Amended soils |
| AY460 | qnrS_1 | QUI | Manure | AY188 | nimE | Other | Amended soils |
| AY553 | msrE | MLSB | Manure | AY421 | aph3-viia | AMI | Amended soils |
| AY559 | catB2 | PHE | Manure | AY430 | ampC_cefa | BL | Amended soils |
| AY56 | ermB_1 | MLSB | Manure | AY431 | blaSME | BL | Amended soils |
| AY568 | tet39 | TET | Manure | AY448 | blaHERA | BL | Amended soils |
| AY75 | lnuA_1 | MLSB | Manure | AY451 | blaLEN | BL | Amended soils |
| AY98 | ampC_1 | BL | Manure | AY469 | fosX | Other | Amended soils |
| AY110 | blaOXA10_1 | BL | Manure | AY541 | vga(A)LC_1 | MLSB | Amended soils |
| AY30 | catB8 | PHE | Manure | AY570 | tet38 | TET | Amended soils |
| AY399 | aac(6)-im | AMI | Manure | AY95 | qnrA | QUI | Amended soils |
| AY4 | aacA/aphD | AMI | Manure | AY484 | bexA/norM | MDR | Vegetable plants |

**TableS2:** List of genes detected in some matrices while absent in others

| id_assay | Gene | Class | High-risk for human-health | Present in | Absent in |
|---|---|---|---|---|---|
| AY181 | vanTG | VAN | no | S1&S3 | S5 |
| AY21 | aadE | AMI | yes | S1&S3 | S5 |
| AY23 | strA | AMI | no | S1&S3 | S5 |
| AY236 | qacEΔ1_3 | Other | no | S1&S3 | S5 |
| AY249 | tet36_1 | TET | no | S1&S3 | S5 |
| AY264 | tetO_2 | TET | yes | S1&S3 | S5 |
| AY269 | tetS | TET | no | S1&S3 | S5 |
| AY274 | tetPB_1 | TET | no | S1&S3 | S5 |
| AY281 | tetM_2 | TET | no | S1&S3 | S5 |
| AY284 | dfrA1_1 | TRI | no | S1&S3 | S5 |
| AY331 | aadA2_3 | AMI | no | S1&S3 | S5 |
| AY368 | tetM_3 | TET | no | S1&S3 | S5 |
| AY423 | aph4-ia | AMI | no | S1&S3 | S5 |
| AY487 | cefa_qacelta | MDR | no | S1&S3 | S5 |
| AY523 | Tn3 | MGE | no | S1&S3 | S5 |
| AY533 | ermB_2 | MLSB | no | S1&S3 | S5 |
| AY534 | ermD | MLSB | no | S1&S3 | S5 |
| AY535 | ermF | MLSB | no | S1&S3 | S5 |
| AY536 | lnuB | MLSB | yes | S1&S3 | S5 |
| AY549 | lnuF | MLSB | no | S1&S3 | S5 |
| AY551 | mefB | MLSB | no | S1&S3 | S5 |
| AY579 | dfrA1 | TRI | yes | S1&S3 | S5 |
| AY580 | dfrA15 | TRI | yes | S1&S3 | S5 |
| AY594 | dfrK | TRI | no | S1&S3 | S5 |
| AY65 | mefA_1 | MLSB | no | S1&S3 | S5 |
| AY549 | lnuF | MLSB | no | S2&S4 | S5 |
| AY142 | ttgB | Other | no | S8&S10 | S9 |
| AY162 | vanHD | VAN | no | S8&S10 | S9 |
| AY204 | sat4 | Other | no | S8&S10 | S9 |
| AY236 | qacEΔ1_3 | Other | no | S8&S10 | S9 |
| AY265 | tetM_1 | TET | no | S8&S10 | S9 |
| AY274 | tetPB_1 | TET | no | S8&S10 | S9 |
| AY276 | tetT | TET | no | S8&S10 | S9 |
| AY331 | aadA2_3 | AMI | no | S8&S10 | S9 |
| AY487 | cefa_qacelta | MDR | no | S8&S10 | S9 |
| AY523 | Tn3 | MGE | no | S8&S10 | S9 |
| AY533 | ermB_2 | MLSB | no | S8&S10 | S9 |
| AY536 | lnuB | MLSB | yes | S8&S10 | S9 |
| AY551 | mefB | MLSB | no | S8&S10 | S9 |
| AY57 | ermT_1 | MLSB | yes | S8&S10 | S9 |
| AY581 | dfra17 | TRI | yes | S8&S10 | S9 |
| AY594 | dfrK | TRI | no | S8&S10 | S9 |
| AY65 | mefA_1 | MLSB | no | S8&S10 | S9 |
| AY83 | ermY | MLSB | no | S8&S10 | S9 |
| AY142 | ttgB | Other | no | S8&S11 | S9 |
| AY162 | vanHD | VAN | no | S8&S11 | S9 |
| AY204 | sat4 | Other | no | S8&S11 | S9 |
| AY218 | qacEΔ1_1 | Other | no | S8&S11 | S9 |
| AY236 | qacEΔ1_3 | Other | no | S8&S11 | S9 |
| AY265 | tetM_1 | TET | no | S8&S11 | S9 |
| AY274 | tetPB_1 | TET | no | S8&S11 | S9 |
| AY276 | tetT | TET | no | S8&S11 | S9 |
| AY331 | aadA2_3 | AMI | no | S8&S11 | S9 |
| AY423 | aph4-ia | AMI | no | S8&S11 | S9 |
| AY487 | cefa_qacelta | MDR | no | S8&S11 | S9 |
| AY523 | Tn3 | MGE | no | S8&S11 | S9 |
| AY533 | ermB_2 | MLSB | no | S8&S11 | S9 |
| AY534 | ermD | MLSB | no | S8&S11 | S9 |
| AY536 | lnuB | MLSB | yes | S8&S11 | S9 |
| AY551 | mefB | MLSB | no | S8&S11 | S9 |
| AY57 | ermT_1 | MLSB | yes | S8&S11 | S9 |
| AY593 | dfrG | TRI | no | S8&S11 | S9 |
| AY594 | dfrK | TRI | no | S8&S11 | S9 |
| AY7 | aphA3_1 | AMI | no | S8&S11 | S9 |
| AY83 | ermY | MLSB | no | S8&S11 | S9 |
| AY142 | ttgB | Other | no | S8&S12 | S9 |
| AY453 | blaBEL | BL | no | S8&S12 | S9 |
| AY537 | lnuC | MLSB | no | S8&S12 | S9 |
| AY551 | mefB | MLSB | no | S8&S12 | S9 |
